# ALMT-independent guard cell R-type anion currents

**DOI:** 10.1101/2022.11.24.517516

**Authors:** Justyna Jaślan, Irene Marten, Liina Jakobson, Triinu Arjus, Rosalia Deeken, Cecilia Sarmiento, Alexis De Angeli, Mikael Bosché, Hannes Kollist, Rainer Hedrich

## Abstract

Plant transpiration is controlled by stomata, with S- and R-type anion channels playing key roles in guard cell action. Arabidopsis mutants lacking the ALMT12/QUAC1 R-type anion channel function in guard cells show only a partial reduction in R-type channel currents. To identify the molecular nature of the remaining R-type anion channel population, patch clamp studies were performed. This R-type current fraction in the *almt12* mutant exhibited the same voltage dependence, susceptibility to ATP block and lacked a chloride permeability as the wildtype. Therefore, we asked whether the R-type anion currents in the ALMT12/QUAC1-free mutant are caused by additional ALMT isoforms. In wildtype guard cells *ALMT12, ALMT13* and *ALMT14* transcripts were detected, whereas only ALMT13 was found expressed in the *almt12* mutant. Substantial R-type anion currents still remained active in the *almt12/13* and *almt12/14* double mutants as well as the *almt12/13/14* triple mutant. This situation, supported by transpiration measurements, suggests that, with the exception of ALMT12, channel species other than ALMTs carry the guard cell R-type anion currents.

## INTRODUCTION

Plants control transpirational water loss via microscopic stomatal pores in the epidermis of leaves. Stomatal movement results from turgor and volume changes in the pore-forming guard cell pairs. The osmotic motor that inflates the guard cells for stomatal opening and deflates them for stomatal closure is powered by changes in potassium salts. The major anionic osmotics that compensate the positive charge of K^+^ are chloride and malate. There is growing evidence that nitrate and sulfate also contribute to stomatal movement (Humble and Hsiao, 1969; Meyer et al., 2010; Malcheska et al., 2017). Patch clamp and microelectrodes studies have shown that voltage-dependent K^+^ channels of the KAT/AKT- and GORK-type mediate guard cell K^+^ uptake and release, respectively (Ache et al., 2000; Szyroki et al., 2001; Ivashikina et al., 2005). For thermodynamic reasons, at physiological negative membrane potentials, anion uptake into guard cells must occur by cotransport with protons. However, the molecular nature of the transporters driving the anions into guard cells is not yet known. In contrast to anion uptake for stomatal opening, guard cell anion release during closure is well understood. Anion efflux is accomplished by two types of anion channels, the S- and R-type (Keller et al., 1989; Schroeder and Hagiwara, 1989; Linder and Raschke, 1992; Negi et al., 2008; Vahisalu et al., 2008; Meyer et al., 2010; Jalakas et al., 2021). In contrast to S-type channels, the R-type anion channels show a pronounced voltage dependence reminiscent of neuronal Na^+^ and Ca^2+^ channels (Hille, 2001; Mumm et al., 2013). S-type anion channels in guard cells are encoded by two genes, *SLAC1* and *SLAH3* (Negi et al., 2008; Vahisalu et al., 2008; Geiger et al., 2011). Both S-type anion channels differ in their relative permeability to chloride versus nitrate (Geiger et al., 2009; Geiger et al., 2011). While SLAC1 is activated by the abscisic acid (ABA) signaling kinase OST1 as well as Ca^2+^-dependent CPK- and CIPK-type kinases, SLAH3 is not regulated by OST1 (Geiger et al., 2009; Geiger et al., 2011; Maierhofer et al., 2014). CPK- and CIPK-dependent activation of SLAH3 requires the presence of extracellular nitrate. In addition, SLAH3 can be activated by physical interaction with the silent but regulatory S-type isoform SLAH1 (Cubero-Font et al., 2016).

In fava bean guard cells R-type anion channels, just like the S-type channels, mediate nitrate and chloride currents. In this plant species the guard cell R-type anion currents are gated by extracellular malate in a process that requires ATP (Hedrich et al., 1990; Hedrich and Marten, 1993; Schmidt and Schroeder, 1994; Dietrich and Hedrich, 1998). In *Arabidopsis thaliana,* R-type currents have been not only identified in guard cells but also in hypocotyl and root epidermal cells (Frachisse et al., 1999; Diatloff et al., 2004; Meyer et al., 2010). In these latter two cell types, the R-type anion currents are found permeable for nitrate, sulfate and chloride.

In search for members of the ALMT family as anion transporter in *Arabidopsis thaliana* guard cells, AtALMT12, a <u>QU</u>ickly activating <u>A</u>nion <u>C</u>hannel (QUAC1), was identified (Meyer et al., 2010). When expressed in *Xenopus laevis* oocytes in the presence of malate, AtQUAC1 gave rise to R-type anion currents. In guard cells of the AtALMT12/QUAC1 loss-of-function mutant, R-type anion currents were reduced but not deficient (Meyer et al., 2010), suggesting that ALMTs other than just ALMT12/QUAC1 are able to form functional R-type anion channels. Here, we ask questions about the molecular fingerprint of R-type anion channel currents. To this end, we systematically knocked out plasma-membrane-expressed ALMT isoforms and compared the guard cell-electrical properties and stomatal transpiration of wildtype and mutant plants.

## RESULTS

### ATP blocks R-type anion channels in guard cells of *Arabidopsis thaliana*

To study the features of R-type currents in guard cells of the AtALMT12-loss-of-function mutant, we performed patch clamp studies in the whole cell configuration similar to those in Meyer et al. (2010). Experiments were conducted in the presence of cytosolic sulfate and ATP (pipette solution) and malate at the extracellular side of the plasma membrane (bath medium). In the voltage-clamp, double voltage pulses were applied to evoke R-type anion currents (Meyer et al., 2010). The recorded R-type inward currents - carried by sulfate efflux – had the same fast activation and deactivation kinetics as guard cell protoplasts of wildtype and the *almt12* mutant (Figs. S1A, 1A). When the steady-state current densities (I/C_m_) were plotted against the clamped membrane voltages (Figs. S1B, 1B), the resulting current/voltage (I/C_m_ (V) curves showed the typical R-type bell shape, peaking around −90 mV. Despite the higher driving force for the anion efflux, the currents declined at more negative voltages and vanished completely negative of −150 mV. Tail currents of wild type and *almt12* (Fig. S1A) revealed the same voltage-dependent open channel probability (Fig. S1C, Table S1). R-type anion channels remained closed at hyperpolarized voltages and opened upon depolarization (Fig. S1C; cf. Mumm et al., 2013), giving rise to the basis of the bell-shaped current/voltage curve (Figs. S1B, 1B). R-type anion currents of *Vicia faba* guard cells and hypocotyl cells of *Arabidopsis thaliana* (Hedrich et al., 1990; Thomine et al., 1997) show a similar bell-shaped voltage dependence (Fig. S1C). AtALMT9, a vacuole localized AtALMT12 homolog, also operates on the basis of a R-type bell-shaped current/voltage curve (Zhang et al., 2014). This vacuolar channel type was found to be ATP-sensitive. To test ATP susceptibility features of R-type anion currents in *Arabidopsis thaliana* guard cells, experiments were performed using cytosolic ATP-free buffers. Wildtype and *almt12* anion currents still exhibited a bell-shaped current/voltage curve (Fig. 1B, C, closed symbols). In the absence of ATP, however, anion peak currents of both wildtype and *almt12* shifted negative by about 70 mV. Due to the increased electrochemical driving force for anion (sulfate) release at this membrane voltage, the peak current amplitudes were more than two times larger than in the presence of ATP. The preserved bell shape of the current/voltage curves in the absence of the adenosine nucleotide (Fig. 1B, C) indicates, that the peculiar voltage dependence is an intrinsic property of wildtype and AtALMT12/QUAC1-free guard cells, which becomes even more pronounced in the presence of ATP (Fig. 1B and C, open symbols). This effect is consistent with the view that ATP occludes the pore of open R-type anion channels, resulting in a voltage-dependent block that becomes progressively stronger with more negative-going membrane voltages, overriding the intrinsic voltagedependent gating.

**Fig. 1.**
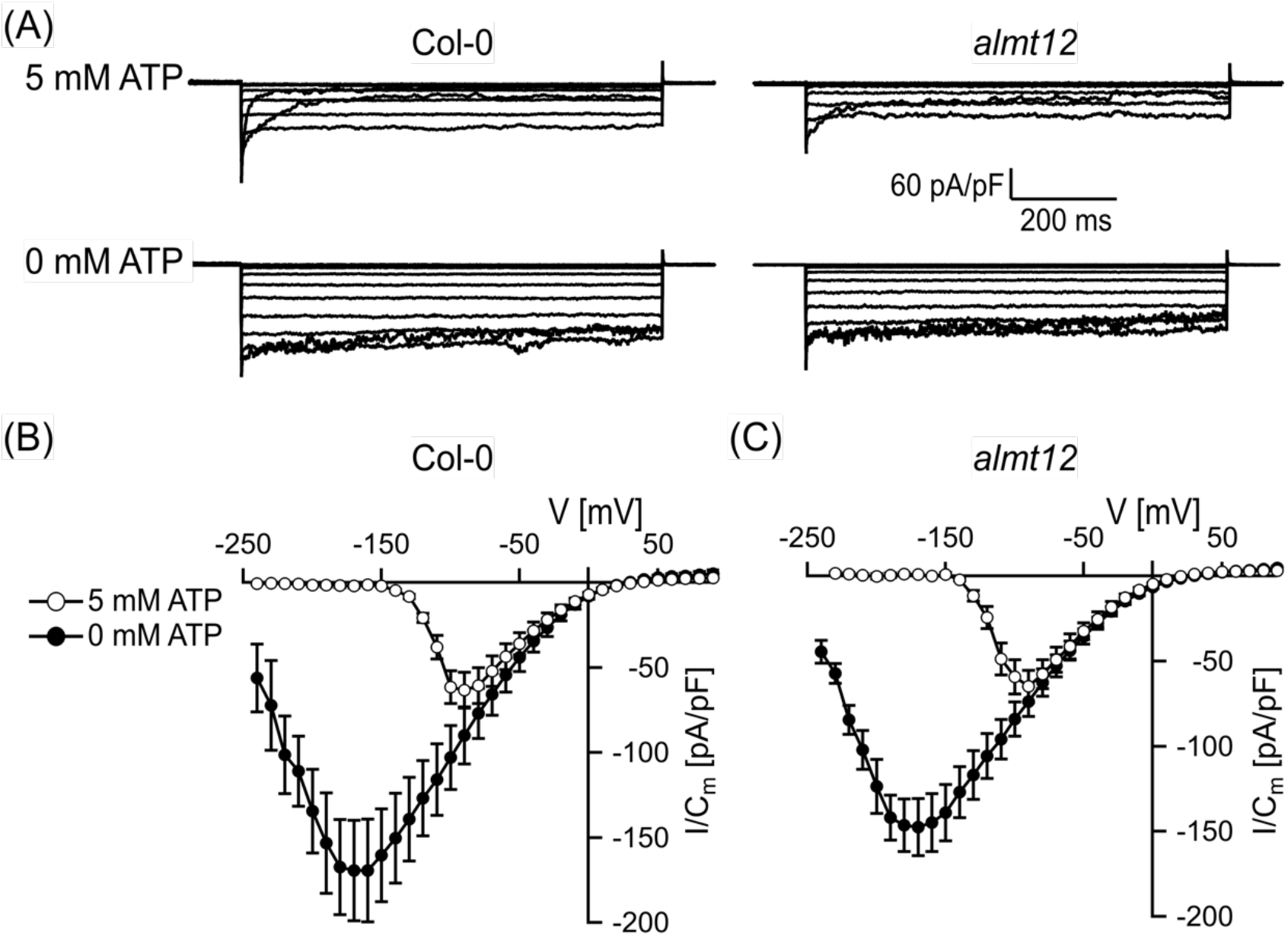
ATP effect on voltage dependence of R-type currents of wildtype and *almt12.* **(A)** Representative voltage-induced current responses. In the presence of ATP, the currents are shown at voltages in the range of −150 and +60 mV in 30-mV steps applied from a holding voltage of −170 mV. In the absence of ATP, the currents are shown at voltages in the range of +70 and −230 mV in 30-mV steps applied from a holding voltage of +70 mV. **(B, C)** Normalized steady-state currents (I/C_m_) of Col-0 (**B**) and *almt12* (**C**) plotted against membrane voltages. Currents were recorded in responses to voltage pulses in the range of +90 mV to −240 mV in 10 mV-decrements. The holding voltage was +70 mV. Number of experiments were n = 5 for Col-0, n = 3 for *almt12* in the presence of ATP (open circles) and n= 6 for Col-0, n = 5 for *almt12* in the absence of ATP (closed circles). Data points represent means ± SEM. Experiments in (**A, B**) were performed in the presence or absence of 5 mM ATP with 195 nM free Ca^2+^ at the cytosolic side of the plasma membrane.

### R-type anion currents in *Arabidopsis thaliana* guard cells are not carried by chloride ions

R-type anion channels in wildtype *Arabidopsis thaliana* hypocotyls and *Vicia faba* guard cells show a permeability for chloride ions (Hedrich and Marten, 1993; Frachisse et al., 1999). When 75 mM sulfate was replaced with equimolar Cl^-^ in the pipette solution supplemented with 1 mM malate, R-type currents were no longer detectable in guard cells of *Arabidopsis thaliana* wildtype and *almt12* mutant plants (Fig. 2, open squares; cf. Fig. S2). Upon supplementing chloride buffer with as low as 1 mM sulfate, small R-type anion currents could be resolved (Fig. 2, open circles). The fact that the peak currents remained at about −90 mV documents that cytosolic sulfate does not act as a gating modifier. This situation further indicates that the AtALMT12/QUAC1 channel component in wildtype as well as the remaining functional R-type channels in the *almt12* mutant are permeable to sulfate but not to chloride.

**Fig. 2.**
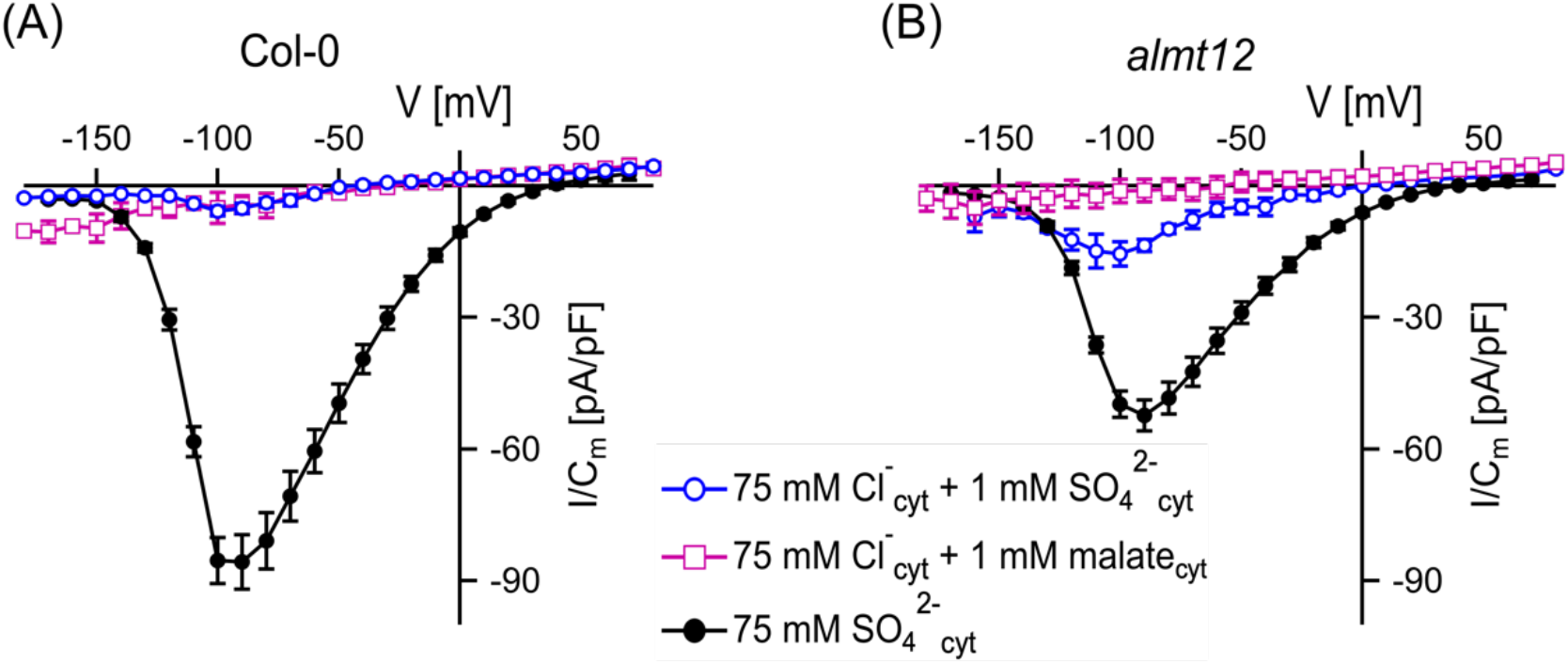
Permeation of R-type anion channels from wildtype (Col-0) and *almt12.* **(A, B)** Normalized whole-cell steady-state currents (I/C_m_) of Col-0 (**A**) and *almt12* (**B**) guard cell protoplasts plotted against the membrane voltages. The holding voltages were −170 mV and −180 mV for current recordings with sulfate- and chloride-based pipette solutions, respectively. Number of experiments were n = 6 for Col-0 (sulfate, closed circles), n = 12 for *almt12* (sulfate, closed circles), n = 3 for Col-0 (chloride/malate, open squares), n = 4 for *almt12* (chloride/malate, open squares), n = 3 for Col-0 and *almt12* (chloride/sulfate, open circles). Data points represent means ± SEM. Experiments were performed in the presence of 195 nM free Ca^2+^ at the cytosolic side of the plasma membrane.

### Guard cells express ALMT13 and ALMT14 in addition to ALMT12/QUAC1

The similar features – in terms of ion permeability, voltage dependence, and ATP susceptibility (Figs. 1, S1, 2) – of the R-type anion currents in guard cells of wildtype and *almt12* mutant plants, led us to speculate that the residual R-type currents in *almt12* originate from other ALMT isoforms. The ALMT family divide into three major clades (Fig. S3). The members ALMT3/4/5/6/9 of clade 2 are localized in the vacuole membrane (Kovermann et al., 2007; Meyer et al., 2011; Takanashi et al., 2016; Eisenach et al., 2017). Because ALMT1/2/11/12, however, are targeted to the plasma membrane (Hoekenga et al., 2006; Meyer et al., 2010; Ligaba et al., 2012; Sasaki et al., 2022), we focused on the ALMTs of clade 1 and 3 (Fig. S3). In quantitative Real-Time PCR experiments with guard-cell protoplast batches of wildtype accession Col-0, we detected transcripts of *ALMT12, ALMT13* and *ALMT14* from clade 3 and ALMT1 from clade 1 (Fig. 3). Of these, *ALMT12* showed the highest expression with a 100-fold higher transcript number. In contrast, extremely few transcripts of ALMT1 were found. We thus focused further analyses on *ALMT13* and *ALMT14* (Fig. 3). *ALMT13* transcripts were detected in guard cells of the single mutant *almt12*, suggesting that the remaining R-type currents in this mutant may be mediated by ALMT13. To test this assumption, we used the double-loss-of-function mutant *almt12/almt13* (Gutermuth et al., 2018) and performed patch clamp studies in the presence of cytosolic sulfate buffers. To our great surprise, however, R-type anion currents of *almt12/almt13* had similar amplitudes and voltage dependencies as those recorded in the *almt12* mutant (Fig. 4). Can these findings be explained by the up-regulated *ALMT14* expression in the *almt12/almt13* mutant background (Fig. 3)? In contrast to the hypothesis, the double mutant *almt12/almt14* (Gutermuth et al., 2018) still showed *almt12-like* R-type currents (Figs. 3, 4, Table S1). Thus, further genetic redundancy could mask the contribution from each of the ALMTs.

**Fig. 3.**
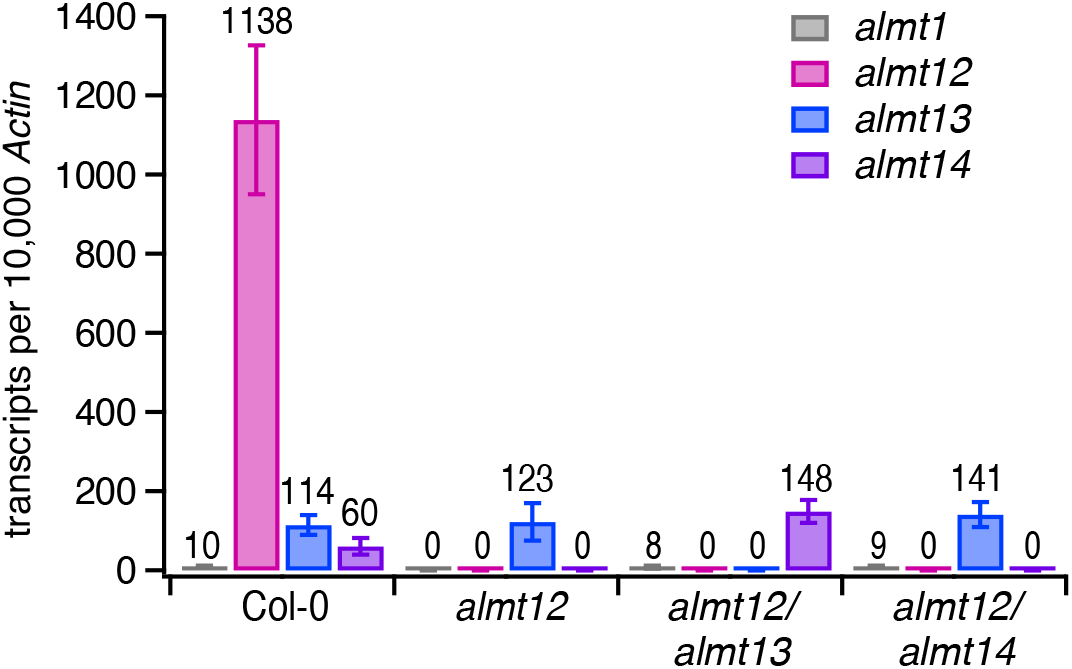
Expression profile of *ALMTs* in *A. thaliana* guard cell protoplasts from wildtype and *almt* mutants. Transcripts of *ALMTs* belonging to clade 1 and III quantified relative to *ACTIN* transcripts. No transcripts of *ALMT2, ALMT7, ALMT8, ALMT10, ALMT11* were detected in all four plant lines. Data represent mean ± SEM. Three biological repetitions were performed. Total number of examined samples was n = 10 for Col-0, n = 7 for *almt12*, n = 7 for *almt12/almt13* and n = 7-8 for *almt12/almt14* loss-of-function mutants. Expression level was quantified for guard cell protoplasts pre-incubated in malate-containing solution. In contrast to *Arabidopsis thaliana* pollen tubes (Gutermuth et al., 2018), the expression of *ALMT11, ALMT12, ALMT13 and ALMT14* in guard cells was not or only slightly affected by malate in the culture media (Fig. S4).

**Fig. 4.**
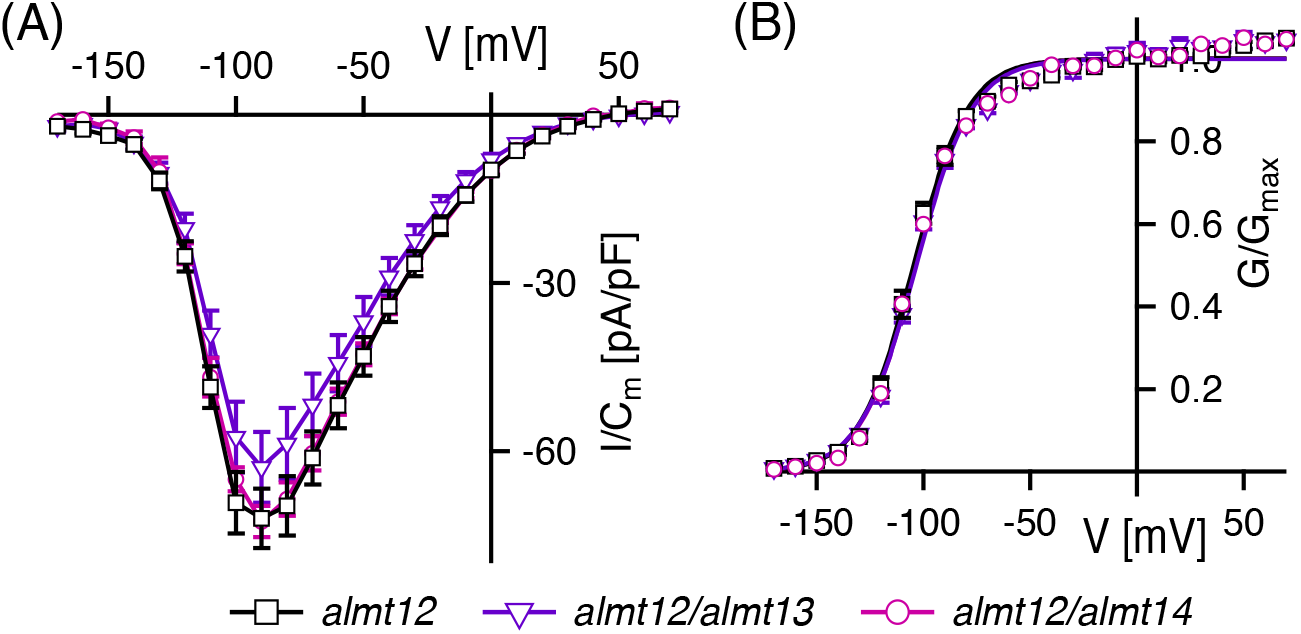
Voltage-dependent R-type currents from single *almt12,* double *almt12/almt13* and *almt12/14* mutants. **(A, B)** Normalized steady-state currents (I/C_m_) and conductances plotted against membrane voltages. The holding voltage was −170 mV. In **(B)** solid lines give the best fits of the data points with the Boltzmann equation. Number of experiments in **(A, B)** were n = 12 for *almt12* (open squares), n = 7 for *almt12/almt13* (open triangles) and n = 3 for *almt12/almt14* (open circles). Data points represent means ± SEM. Experiments were conducted with 1.85 μM free Ca^2+^ at the cytosolic side of the plasma membrane.

### Generation of *almt12/13/14* triple mutant lines

To decipher whether ALMT13 and ALMT14, like ALMT12, could function as active R-type anion channels in guard cells, a triple mutant lacking the function of all three ALMTs was generated. We used CRISPR-Cas9 technology to inactivate *ALMT13* and *ALMT14* in the background of the *ALMT12* T-DNA insertion line, because *ALMT13* and *ALMT14* are localized next to each other on chromosome 5 (At5g46600 and At5g46610, respectively) which precludes the possibility of crossing the T-DNA-insertion lines. Three different CRISPR-induced lines were created in the background of *almt12-2* (SM_3_1713) and verified by Sanger sequencing of the genomic and coding DNA fragments (Table 1). Line *almt12/almt13/almt14#08* carries (i) in *ALMT12* a transposon insert, (ii) in *ALMT13* six bp deletion at positions 125-130 (ATGTGG) from ATG leading to amino acid changes (41WNVGK45→41WRK43) and (iii) in *ALMT14* 14 bp deletion at positions 54-67 from ATG together with an additional single nucleotide polymorphism (G32A, G11E) leading to an early stop codon after 21 amino acid residues. Line *almt12almt13almt14 #11* carries (i) in *ALMT12* a transposon insert, (ii) in *ALMT13* an additional +G at position 129 from ATG onwards, leading to an early stop codon after 92 amino acid residues and (iii) in *ALMT14* an additional +T at position 68 from ATG onwards, leading to early stop codon after 26 amino acid residues. Line *almt12/almt13/almt14#19* carries (i) in *ALMT12* a transposon insert, (ii) in *ALMT13* an additional +G at position 129 from ATG onwards, leading to an early stop codon after 92 amino acid residues and (iii) in *ALMT14* an additional +A at position 67 from the first ATG onwards, resulting in an early stop codon after 26 amino acid residues. A multiple sequence alignment depicted the mutagenesis sites in the coding sequences induced by the CRISPR-Cas9 system in the aforementioned triple mutant lines (Fig. S5). The cDNAs of *ALMT13* and *ALMT14* were sequenced in all *almt* triple mutant lines in the range of 90 bp upstream of the ATG start codon up to the stop codon. We verified that the cDNAs were similar to the native sequences with the exceptions of CRISPR-Cas9-induced mutated sites mentioned above. No other rearrangements such as exon skipping, or alternative splice sites were encountered. Thus, we conclude that the abovementioned *almt* triple mutant lines *almt12/almt13/almt14#08, almt12/almt13/almt14#11* and *almt12/almt13/almt14#19* express the above-described mutated versions of *ALMT12, ALMT13* and *ALMT14* and that these proteins are therefore non-functional in these triple mutants.

**Table 1.** CRISPR-Cas-induced mutations were generated in the first exons of *AtALMT13* and *AtALMT14* in the background of *almt12-2.*

| Line | *AtALMT12* | *AtALMT13* | *AtALMT14* |
| --- | --- | --- | --- |
| *almt12-2* (SM_3_1713) | transposon insert | WT | WT |
| *almt12/almt13/almt14#08* | transposon insert | -6 bp | -14 bp + 1 SNP |
| *almt12/almt13/almt14 #11* | transposon insert | +G | +T |
| *almt12/almt13/almt14#19* | transposon insert | +G | +A |

### Loss of ALMT13 and ALMT14 function affect neither guard cell R-type anion currents nor stomatal transpiration

To finally answer questions regarding the ALMT nature of R-type currents, patch-clamp recordings were performed on guard cell protoplasts from the triple *almt12/13/14#19* mutant. Under the same conditions as described above for the double *almt* mutants, voltage stimulation of the triple *almt* mutant evoked pronounced R-type anion currents (Fig. 6A, B) of similar voltage dependence as guard cells from the single/double mutants *almt12*, *almt12/13* and *almt12/14* (Figs. 4B, 6C, Table S1). The electrical fingerprint of the triple mutant demonstrates that ALMT13 and ALMT14 do not represent major channel components of R-type anion currents in the single and double *almt* mutants and therefore, unlike ALMT12, do not contribute to the currents in wildtype.

Meyer et al. (2010) showed that the ALMT12/QUAC1-loss-of-function mutant was impaired in CO_2_-induced stomatal closure. In analogy, we also performed gas exchange measurements, using transpiratory water loss as a proxy for stomatal action (Fig. 5). To trigger stomatal responses, Arabidopsis wildtype plants and single/double/triple *almt* mutants *(almt12, almt13, almt14, almt12/13, almt12/14, almt12/13/14)* were challenged with changes in atmospheric CO_2_ levels (Fig. 5B). A decrease in [CO_2_] from 400 to 100 ppm resulted in an equally strong and rapid stomatal opening of wildtype, *almt12 and almt12/13/14,* while a subsequent rise from 100 to 400 and 800 ppm caused stomatal closure in wildtype. This CO_2_-triggered response of stomatal guard cells, however, was much less pronounced in all three triple mutant lines and the single *almt12* mutant to the same extent (Fig. 5B). Furthermore, the stomatal conductance of the double mutants *almt12/13* and *almt12/14* was impaired in a similar manner to *almt12* (Fig. S6). Stomata of the single *almt13* and *almt14* mutants, however, responded to higher CO_2_ levels just as those of wildtype plants (Fig. S6). These results show that loss of ALMT13 or ALMT14 function alone did not affect stomatal closure, nor did it further enhance the disruption of stomatal closure in *almt12.* Accordingly, ALMT12 but not ALMT13 and ALMT14 is essential for CO_2_-induced stomatal closure.

**Fig. 5.**
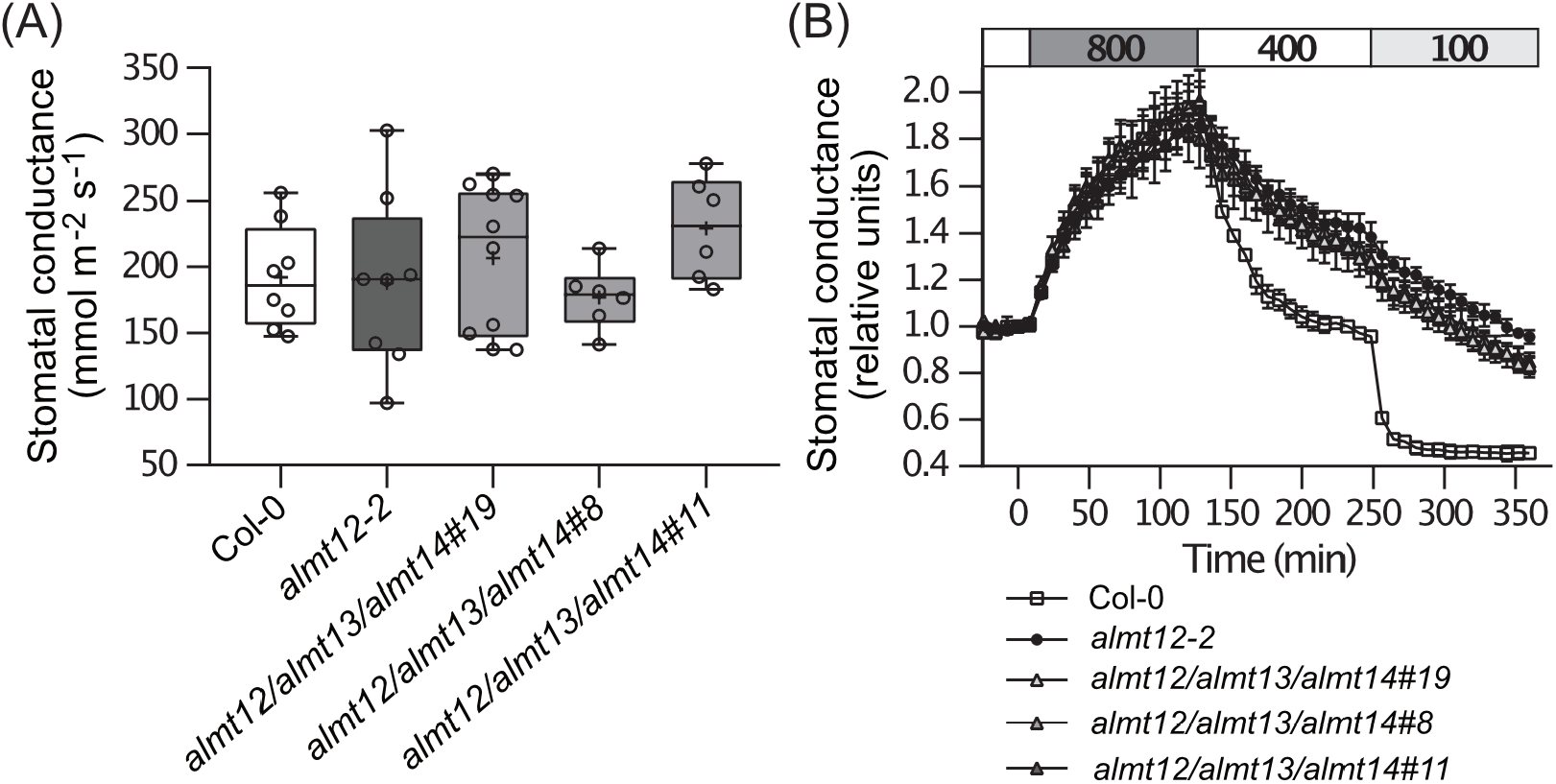
Gas exchange of different triple *almt12/almt13/almt14* mutant lines. **(A)** Box plot of steady-state stomatal conductance values (mmol m^-2^ s^-1^) of every measured plant at a CO_2_ level of 400 ppm. Boxes represent 25-75 % quartiles and the horizontal line in the box is the median, whiskers indicate the smallest and largest values, points mark individual plant stomatal conductance values. There were no statistically significant differences between the groups (One-way ANOVA with Tukey *post hoc* test). **(B)** Time-resolved patterns of normalized stomatal conductances during changing CO_2_ levels from 400 ppm to 100 ppm, then elevation from 100 ppm to 400 ppm and from 400 ppm to 800 ppm. Mean normalized stomatal conductance ± SEM are given.

## DISCUSSION

R-type anion currents in guard cells are only partially mediated by AtALMT12/QUAC1 (Fig. S1; Meyer et al., 2010). The remaining R-type currents in the *almt12* mutant share major features with wildtype R-type currents such as fast activation/deactivation, voltage dependence and ATP susceptibility, as well as a dominant sulfate permeability (Fig. 1, 2, S1). Nevertheless, they are most likely generated not by other ALMT species but by other channel specie(s), because (i) R-type currents did not disappear with concomitant loss of ALMT12/13/14 function, and (ii) CO_2_-induced stomatal closure depended on ALMT12 but not on ALMT13 or ALMT14 (Figs. 5, 6).

**Fig. 6.**
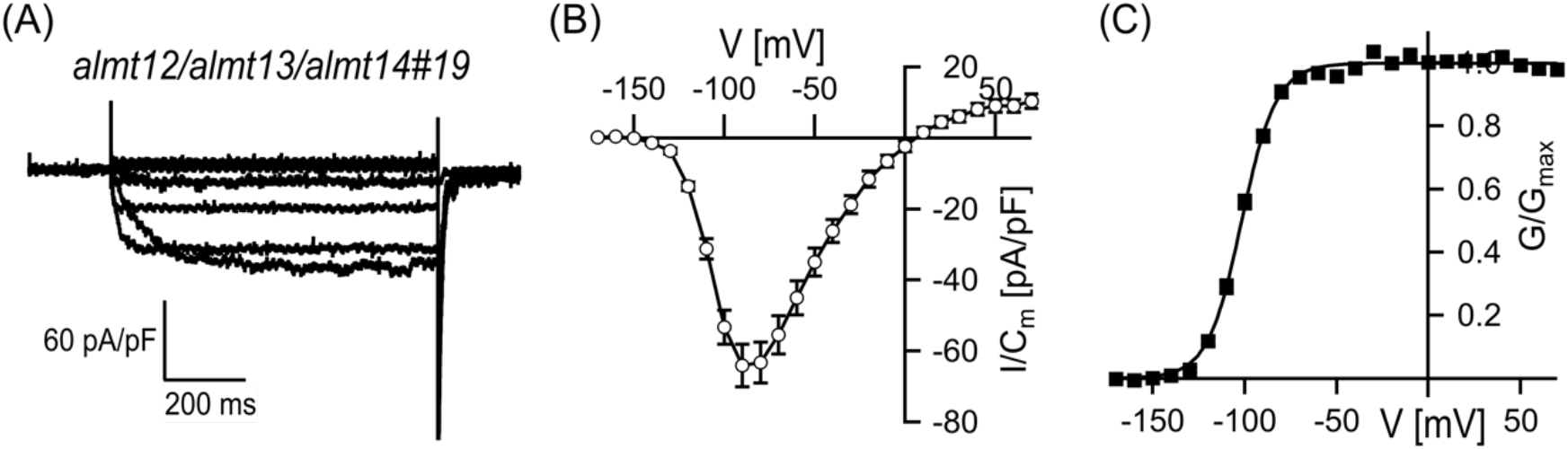
Voltage-dependent R-type currents from triple *almt12/almt13/almt14* mutants. **(A)** Representative whole-cell currents shown at voltages in the range of −150 and +60 mV in 30-mV steps. **(B, C)** Normalized steady-state currents (I/C_m_) and conductances (G/G_max_) plotted against membrane voltages. Number of experiments were n=14. Data points represent means ± SEM. Experiments were conducted with 1.85 μM free Ca^2+^ at the cytosolic side of the plasma membrane. (**A-C**) The holding voltage was −170 mV.

CryoEM analyses with the AtALMT12 homolog from soybean (GmALMT12) identified several positively charged amino acids decorating the ion conductive pore of R-type channel component (Qin et al., 2022). In line with these positive charges forming the selectivity filter, AtALMT12, when expressed in *Xenopus laevis* oocytes, mediates only anion currents (Meyer et al., 2010; Sasaki et al., 2010). In the latter heterologous expression system, AtALMT12/QUAC1 is characterized by the typical bell-shaped R-type I/V curve and is highly permeable to the dicarboxylate anions fumarate and malate (Meyer et al., 2010; Sasaki et al., 2022). Although ALMT12/QUAC1 is predominately expressed in *A. thaliana* guard cells (Fig. 3), the guard cell R-type currents are not carried by chloride ions, even in the presence of activating malate on both sides of the membrane (Fig. 2, S2). In contrast to *A. thaliana* guard cells (Fig. 2A, S1), R-type channels in *A. thaliana* hypocotyls and *Vicia faba* guard cells have major conductance for chloride and nitrate but are weakly permeable to malate (Hedrich and Marten, 1993; Frachisse et al., 1999). ALMT12 channels alone could thus not explain the permeability for monovalent anions in guard cells of species such as *V. faba* or in hypocotyl cells of *A. thaliana.*

Malate is not only a permeating substrate but also acts as a gating modulator of AtALMT12, leading to a shift in the voltage activation threshold of AtALMT12 to more negative membrane voltages and in turn channel activation (Meyer et al., 2010; Imes et al., 2013). The stimulatory effect of malate on GmALMT12 channel activity appears to be due to the interaction of the dicarboxylate with pore residues accessible from both sides of the membrane (Qin et al., 2022). Accordingly, ALMT12 alone also could not explain the malate modulation of R-type channel gating in *V. faba,* where malate acts only from the outside (Hedrich and Marten, 1993).

Moreover, R-type channels of *A. thaliana* and *V. faba* respond differently to ATP. In *A. thaliana* guard cells (Fig. 1) and hypocotyl cells (Thomine et al., 1997; Colcombet et al., 2001), ATP inhibits the R-type anion currents in a voltage-dependent manner, shifting the voltage activation threshold to more positive voltages. In this respect, the inhibitory ATP effect on the plasma membrane R-type currents of *A. thaliana* resembles that on the vacuolar anion currents mediated by AtALMT9 (Zhang et al., 2014). The ATP block on AtALMT9 is caused by an ATP interaction with a conserved lysine at position 193 in the AtALMT9 channel pore (Zhang et al., 2014). Since AtALMT12 contains a lysine at a homologous site (Lys154), the ATP susceptibility of R-type currents in *A. thaliana* wildtype guard cells could partially reflect an ALMT9-like ATP block of the AtALMT12 pore. The remaining currents in the *almt12* mutant, however, could not be associated with an ALMT conductance (Fig. 6) and continued to respond to ATP in the same manner as the wildtype currents (Fig. 1). Therefore, the observed ATP effect on R-type currents of *A. thaliana* guard cells could be related additionally or even exclusively to an unknown R-type anion channel component. In contrast to *A. thaliana*, ATP functions as an activator of R-type anion channels in *V. faba* guard cells without changing their voltage-dependent gating behavior (Hedrich et al., 1990). Thus, cell type- and species-specific differences and similarities in certain R-type current features appear to exist. It now remains to be elucidated which channel type(s) contribute(s) to the R-type anion currents in these two species. Taken together, our findings relaunch the race to identify the overall molecular nature of the R-type anion conductances in different species and cell types.

## ACKNOWLEDGEMENTS

The work was funded by the German DFG Koselleck award (HE 1640/42-1) to RH and by the Estonian Research Council (PRG433) and European Regional Development Fund (Centre of Excellence in Molecular Cell Engineering, CEMCE) to HK, and by a CNRS ATIP-AVENIR and by the French Research Agency (ANR-PARASOL) to ADA. The group of RH thank Joachim Rothenhöfer for plant cultivation and Brigitte Neumann for technical assistance. We are grateful to Dr. Pirko Jalakas for technical assistance.

## AUTHOR CONTRIBUTION

IM, RH designed the electrophysiological research work; JJ performed the patch clamp experiments and analyzed the current recordings; ADA analyzed part of the patch-clamp currents; RD designed and JJ performed the qRT-PCR experiments; JJ and RD analysed the qRT-PCR data. HK designed the gas exchange experiments and together with CS the generation of the triple *almt* mutant; MB isolated single and double mutants; LJ created the triple *almt* mutants; TA performed and analyzed the gas exchange experiments. IM and RH wrote the manuscript, and all authors revised the manuscript.

## MATERIAL AND METHODS

### Plant Material and Growth Condition

*Arabidopsis thaliana* wild type plants (Col-0) and all different ALMT-loss-of-function mutants *(almt12, almt12/almt13, almt12/almt14, almt12/almt13/almt14*) were cultivated in soil in a growth chamber under short day condition (8/16 h light/dark regime) with a photon flux density of about 150 μmol·m-^2^·s-^1^ at 60% relative humidity. The temperature in the growth chamber was kept at 22°C and 16°C during the day and night, respectively.

### Selection of *almt12, almt12/almt13* and *almt12/almt14* T-DNA Mutant Lines

The *atalmt12* single mutant line (SM_3_38592) was genotyped with same AtALMT12- and T-DNA-specific primers as previously described by (Meyer et al., 2010). Double-knockout mutant lines *almt12/almt13* and *almt12/almt14* were created by crossing *atalmt12-1* (SM_3_38592) with either *atalmt13* (SAIL_517_F03) or *atalmt14* (SALK_046375C) (Gutermuth et al., 2018). For PCR genotyping of the double-knockout mutants, same ALMT12 primers were used as for the single *almt12* mutant while following gene- and T-DNA-specific primers were used for ALMT13 and ALMT14: ALMT13fwd (5’-GGCGGATTTAGAATCCAAAAC-3’), ALMT13rev (5’-TAATAAAGCGGAATTGGAACG-3’), SAIL-tDNA (5’-TTCATAACC AATCTCGA TACAC-3’), ALMT14fwd (5’-TTGGAAACAGTGCTATTTGGG-3’), ALMT14rev (5’-TGAGGCCGAAAGTATAAAAAGC-3’) and SALK-Lba (5’-TGGTTCACGTAGTGGGCCATCG-3’).

### CRISPR-Cas9-induced mutagenesis

For site-directed mutagenesis with CRISPR-Cas9 system, the binary vector pHEE401E (Addgene) with two gRNAs was used. The plasmid expresses an egg-cell-specific promoter-controlled cassette of 3xFLAG-NLS-zCas9-NLS together with hygromycin resistance gene. *ALMT13* (At5g46600) gRNA (GAATCTATGGAATGTGGGAAAGG) targeted the cut site 130 bp from ATG in the first exon. *ALMT14* (At5g46610) gRNA (TACTAAAAACATGAAGACAAAGG) targeted the cut site 67 bp from ATG also in the first exon. *atalmt12-2* plants (SM_3_1713) were transformed with *Agrobacterium tumefaciens* strain C58C1(pTiB6S3ΔT-DNA) carrying the binary vector with the floral dip method (Clough and Bent, 1998).

### Selection of T-DNA-free and hygromycin-sensitive CRISPR plants

For selecting for hygromycin-sensitive T2 seeds, the seeds were germinated on hygromycin-free MS plates for 7-8 days. Then, the seedlings were transferred to hygromycin-containing MS plates (25-50 μg/ml). 1-2 days later the seedlings with no primary root growth were rescued and transferred to soil.

### gDNA and cDNA Sequencing of *ALMT13* and *ALMT14*

Genomic DNA was extracted either from leaves of 5-weeks old plants or from 7-days-old plate-grown seedlings (in the case of homozygous lines). DNA extraction buffer (100 mM TRIS, pH=8.0; 50 mM EDTA; 500 mM NaCl) together with the plant tissue was ground by micro pestle, then centrifuged for 5 min at maximum speed and precipitated with isopropanol. For sequencing *ALMT13* cDNA, RNA was extracted from leaves with the E.Z.N.A.® Plant RNA Kit (Omega Bio-tek) according to the manufacturer’s protocol. For sequencing *ALMT14* cDNA, RNA was extracted from 50-80 mg of surface-sterilised (70% EtOH, 0.1% Triton X-100) and 3-days germinated seeds with RNeasy Mini kit (Qiagen) according to the manufacturer’s protocol. Extracted RNA was thereafter treated with DNase I (Thermo Fischer Scientific) and cDNA was synthesized with Maxima H Minus reverse transcriptase (Thermo Fischer Scientific). PCR and RT-PCR were carried out with Phusion High-Fidelity DNA polymerase (Thermo Fischer Scientific) using the following PCR programme: 98°C for 3 min; 40 cycles of 98°C for 10 s, 63-64.7°C for 30 s, 72°C 90 s; 72°C for 10 min. Primers used in the PCRs are depicted in Table S2.

### Quantitative Real-Time PCR (qRT PCR) with Guard Cell Protoplast RNA

The number of transcripts were determined for guard cell protoplasts enzymatically isolated from leaves of five-week-old plants. For protoplast isolation, the same protocol was used as for patch clamp experiments (see below). After enrichment of the protoplasts, these were stored for 1 h at room temperature in washing solution in the absence and presence of 20 mM malate, as indicated. Then, the protoplast samples were centrifuged again at 69 *g* and 4°C for 14 min, and the supernatant was discarded. RNA was isolated from the plant material using the RNeasy Plant Mini Kit (www.quiagen.com) according to the manufacturer’s protocol. For cDNA synthesis total RNA was extracted as described by Logemann et al. (1987). Using 2.5 μg of total RNA, cDNA was synthesized according to Biemelt et al. (2004). For quantification of the number of *ALMT* transcripts quantitative real-time PCR (qRT PCR) was performed as previously described by Geiger et al. (2011). Transcript were normalized to 10.000 molecules of *AtACTIN2/ACTIN8* using standard curves calculated for the individual PCR products. Oligonucleotides used for qRT PCR are listed in Table S3.

### Gas exchange experiments

*Arabidopsis thaliana* wildtype Col-0, mutant lines *almt12-2* and CRISPR/Cas9-generated triple *almt12/almt13/almt14* mutant lines #19, #8 and #11 were sown in custom gas exchange pots with 4:2:3 peat:vermiculite:water growth medium and grown in controlled-environment growth cabinets (AR-66LX; Percival Scientific) at 10/14 photoperiod with 250 μmol m^-2^ s^-1^ white light, 70% relative humidity, day-time temperature of 23°C and night temperature of 18°C. Plants were measured at 24-31 days of age. Whole plant rosette gas-exchange measurements were conducted with custom-built gas-exchange device (Kollist et al., 2007; Plant Invent Ltd.). Plant was inserted into measurement cuvette and allowed to stabilize for ~2 hours at 250 μmol m^-2^ s^-1^ light intensity, ~70% relative air humidity, 23°C air temperature and 400 ppm CO_2_ concentration. For testing stomatal responses in response to various CO_2_ levels, the CO_2_ concentration was adjusted from 400 ppm to 100 ppm for 2 hours. From there, CO_2_ levels were restored to 400 ppm for two hours before applying 800 ppm concentration of CO_2_ for the next two hours. For testing stomatal responses to low air humidity, air with 20% relative humidity was applied for 56 minutes and thereafter 70% relative air humidity was restored. Statistical analysis for Fig. 5A was carried out with Past326b.

### Patch Clamp Experiments on Guard Cell Protoplasts

For patch clamp recordings, guard cell protoplasts were isolated as previously described in (Geiger et al., 2010). In brief, the lower epidermis was peeled off from leaves of 5-6 week-old plants and incubated in enzyme solution for 16 h at 18°C. Then, the suspension was washed through a 50 μm nylon mesh using a wash solution that was composed of 20 mM CaCl_2_, 5 mM MES, pH 5.6/Tris, 400 mOsmol kg^-1^ adjusted with D-sorbitol. After centrifugation at 69 *g* and 4°C for 14 min and removal of the supernatant, the protoplast-enriched pellet was stored on ice until protoplasts were used for patch clamp measurements. For this, the whole-cell configuration was established on protoplasts using glass microelectrodes with a typical patch-pipette resistance of about 3-5 Mohm prepared from GB150T-8P glass capillaries (Science Product). Patch-clamp experiments were carried out using standard bath solution (2 mM MgCl_2_, 0.5 mM LaCl3, 10 mM MES, 20 mM CaMalate, pH 5.6/Tris) and standard patch-pipette solution (75 mM Cs2SO_4_, 2 mM MgCl_2_, 5 mM Mg-ATP, 10 mM HEPES, pH 7.1/Tris). The free Ca^2+^ concentration of the pipette medium was additionally adjusted to either 195 nM or 1.85 μM by adding certain quantities of CaCl_2_ and EGTA as calculated with WEBMAXC standard (https://somapp.ucdmc.ucdavis.edu/pharmacology/bers/maxchelator/webmaxc/webmaxcS.htm). To study chloride permeation, chloride-based pipette medium was used, containing 75 mM CsCl instead of 75 mM Cs2SO_4_, and was eventually supplemented with 1 mM Cs2SO_4_ or 1 mM malate as indicated. To examine the ATP effect on channel function, some patch-clamp experiments were performed with ATP-free pipette solutions. The osmolality of the bath and pipette solutions was adjusted to 400 and 440 mOsmol kg^-1^ with D-sorbitol, respectively.

For macroscopic current recordings, two different voltage-pulse protocols were applied to the plasma membrane. From a holding voltage of −170 mV or −180 mV, 800-ms-lasting voltage pulses were applied in the range of −170 mV to +70 mV in 10-mV increments. When the plasma membrane was clamped to a holding voltage of +70 mV, the 800-ms-lasting voltage pulses were applied in the range of +90 mV to −240 mV in 10-mV decrements. The clamped voltages were corrected off-line for the liquid junction potential as described in Neher (1992). Seven minutes after the whole-cell configuration was established, membrane currents were recorded at a sampling rate of 100 μs and low-pass filtered at 1 kHz. For comparison of different guard cell protoplasts, steady-state currents were normalized to the membrane capacitance (C_m_) as a measure for the size of each guard cell protoplast. As a measure for the voltage-dependent relative open-channel probability, conductance/voltage curves (G/G_max_(V)) were quantified from instantaneous tail current amplitudes recorded at −170 mV after each prepulse-voltage ranging from −170 mV to +70 mV. After normalization to maximum predicted tail current density, values were plotted against the corresponding prepulse voltages and fitted with a Boltzmann equation providing the midpoint voltage (V_1/2_) and the apparent equivalent gating charge (z).

## REFERENCES

Ache, P., Becker, D., Ivashikina, N., Dietrich, P., Roelfsema, M.R., and Hedrich, R. (2000). GORK, a delayed outward rectifier expressed in guard cells of Arabidopsis thaliana, is a K(+)-selective, K(+)-sensing ion channel. FEBS Lett 486:93–98.

Biemelt, S., Tschiersch, H., and Sonnewald, U. (2004). Impact of altered gibberellin metabolism on biomass accumulation, lignin biosynthesis, and photosynthesis in transgenic tobacco plants. Plant Physiol 135:254–265.

Clough, S.J., and Bent, A.F. (1998). Floral dip: a simplified method for Agrobacterium-mediated transformation of Arabidopsis thaliana. Plant J 16:735–743.

Colcombet, J., Thomine, S., Guern, J., Frachisse, J.M., and Barbier-Brygoo, H. (2001). Nucleotides provide a voltage-sensitive gate for the rapid anion channel of arabidopsis hypocotyl cells. J Biol Chem 276:36139–36145.

Cubero-Font, P., Maierhofer, T., Jaslan, J., Rosales, M.A., Espartero, J., Diaz-Rueda, P., Muller, H.M., Hurter, A.L., Al-Rasheid, K.A., Marten, I., et al. (2016). Silent S-Type Anion Channel Subunit SLAH1 Gates SLAH3 Open for Chloride Root-to-Shoot Translocation. Curr Biol 26:2213–2220.

Diatloff, E., Roberts, M., Sanders, D., and Roberts, S.K. (2004). Characterization of anion channels in the plasma membrane of Arabidopsis epidermal root cells and the identification of a citrate-permeable channel induced by phosphate starvation. Plant Physiol 136:4136–4149.

Dietrich, P., and Hedrich, R. (1998). Anions permeate and gate GCAC1, a voltage-dependent guard cell anion channel. Plant Journal 15:479–487.

Eisenach, C., Baetz, U., Huck, N.V., Zhang, J., De Angeli, A., Beckers, G.J.M., and Martinoia, E. (2017). ABA-Induced Stomatal Closure Involves ALMT4, a Phosphorylation-Dependent Vacuolar Anion Channel of Arabidopsis. Plant Cell 29:2552–2569.

Frachisse, J.M., Thomine, S., Colcombet, J., Guern, J., and Barbier-Brygoo, H. (1999). Sulfate is both a substrate and an activator of the voltage-dependent anion channel of Arabidopsis hypocotyl cells. Plant Physiol 121:253–262.

Geiger, D., Maierhofer, T., Al-Rasheid, K.A., Scherzer, S., Mumm, P., Liese, A., Ache, P., Wellmann, C., Marten, I., Grill, E., et al. (2011). Stomatal closure by fast abscisic acid signaling is mediated by the guard cell anion channel SLAH3 and the receptor RCAR1. Sci Signal 4:ra32.

Geiger, D., Scherzer, S., Mumm, P., Marten, I., Ache, P., Matschi, S., Liese, A., Wellmann, C., Al-Rasheid, K.A., Grill, E., et al. (2010). Guard cell anion channel SLAC1 is regulated by CDPK protein kinases with distinct Ca2+ affinities. Proc Natl Acad Sci U S A 107:8023–8028.

Geiger, D., Scherzer, S., Mumm, P., Stange, A., Marten, I., Bauer, H., Ache, P., Matschi, S., Liese, A., Al-Rasheid, K.A., et al. (2009). Activity of guard cell anion channel SLAC1 is controlled by drought-stress signaling kinase-phosphatase pair. Proc Natl Acad Sci U S A 106:21425–21430.

Gutermuth, T., Herbell, S., Lassig, R., Brosche, M., Romeis, T., Feijo, J.A., Hedrich, R., and Konrad, K.R. (2018). Tip-localized Ca(2+) -permeable channels control pollen tube growth via kinase-dependent R- and S-type anion channel regulation. New Phytol 218:1089–1105.

Hedrich, R., Busch, H., and Raschke, K. (1990). Ca2+ and nucleotide dependent regulation of voltage dependent anion channels in the plasma membrane of guard cells. EMBO J 9:3889–3892.

Hedrich, R., and Marten, I. (1993). Malate-induced feedback regulation of plasma membrane anion channels could provide a CO2 sensor to guard cells. EMBO J 12:897–901.

Hille, B. (2001). Ionic channels of excitable membranes. Sunderland, Mass: Sinauer Associates.

Hoekenga, O.A., Maron, L.G., Pineros, M.A., Cancado, G.M., Shaff, J., Kobayashi, Y., Ryan, P.R., Dong, B., Delhaize, E., Sasaki, T., et al. (2006). AtALMT1, which encodes a malate transporter, is identified as one of several genes critical for aluminum tolerance in Arabidopsis. Proc Natl Acad Sci U S A 103:9738–9743.

Humble, G.D., and Hsiao, T.C. (1969). Specific requirement of potassium for light-activated opening of stomata in epidermal strips. Plant Physiol 44:230–234.

Imes, D., Mumm, P., Bohm, J., Al-Rasheid, K.A., Marten, I., Geiger, D., and Hedrich, R. (2013). Open stomata 1 (OST1) kinase controls R-type anion channel QUAC1 in Arabidopsis guard cells. Plant J 74:372–382.

Ivashikina, N., Deeken, R., Fischer, S., Ache, P., and Hedrich, R. (2005). AKT2/3 subunits render guard cell K+channels Ca2+ sensitive. J Gen Physiol 125:483–492.

Jalakas, P., Nuhkat, M., Vahisalu, T., Merilo, E., Brosche, M., and Kollist, H. (2021). Combined action of guard cell plasma membrane rapid-and slow-type anion channels in stomatal regulation. Plant Physiol 187:2126–2133.

Keller, B.U., Hedrich, R., and Raschke, K. (1989). Voltage-Dependent Anion Channels in the Plasma-Membrane of Guard-Cells. Nature 341:450–453.

Kollist, T., Moldau, H., Rasulov, B., Oja, V., Ramma, H., Huve, K., Jaspers, P., Kangasjarvi, J., and Kollist, H. (2007). A novel device detects a rapid ozone-induced transient stomatal closure in intact Arabidopsis and its absence in abi2 mutant. Physiol Plantarum 129:796–803.

Kovermann, P., Meyer, S., Hortensteiner, S., Picco, C., Scholz-Starke, J., Ravera, S., Lee, Y., and Martinoia, E. (2007). The Arabidopsis vacuolar malate channel is a member of the ALMT family. Plant J 52:1169–1180.

Ligaba, A., Maron, L., Shaff, J., Kochian, L., and Pineros, M. (2012). Maize ZmALMT2 is a root anion transporter that mediates constitutive root malate efflux. Plant Cell Environ 35:1185–1200.

Linder, B., and Raschke, K. (1992). A slow anion channel in guard cells, activating at large hyperpolarization, may be principal for stomatal closing. FEBS Lett 313:27–30.

Logemann, J., Schell, J., and Willmitzer, L. (1987). Improved method for the isolation of RNA from plant tissues. Anal Biochem 163:16–20.

Maierhofer, T., Diekmann, M., Offenborn, J.N., Lind, C., Bauer, H., Hashimoto, K., Ka, S.A.-R., Luan, S., Kudla, J., Geiger, D., et al. (2014). Site- and kinase-specific phosphorylation-mediated activation of SLAC1, a guard cell anion channel stimulated by abscisic acid. Sci Signal 7:ra86.

Malcheska, F., Ahmad, A., Batool, S., Muller, H.M., Ludwig-Muller, J., Kreuzwieser, J., Randewig, D., Hansch, R., Mendel, R.R., Hell, R., et al. (2017). Drought-Enhanced Xylem Sap Sulfate Closes Stomata by Affecting ALMT12 and Guard Cell ABA Synthesis. Plant Physiol 174:798–814.

Meyer, S., Mumm, P., Imes, D., Endler, A., Weder, B., Al-Rasheid, K.A., Geiger, D., Marten, I., Martinoia, E., and Hedrich, R. (2010). AtALMT12 represents an R-type anion channel required for stomatal movement in Arabidopsis guard cells. Plant J 63:1054–1062.

Meyer, S., Scholz-Starke, J., De Angeli, A., Kovermann, P., Burla, B., Gambale, F., and Martinoia, E. (2011). Malate transport by the vacuolar AtALMT6 channel in guard cells is subject to multiple regulation. Plant J 67:247–257.

Mumm, P., Imes, D., Martinoia, E., Al-Rasheid, K.A., Geiger, D., Marten, I., and Hedrich, R. (2013). C-terminus-mediated voltage gating of Arabidopsis guard cell anion channel QUAC1. Mol Plant 6:1550–1563.

Negi, J., Matsuda, O., Nagasawa, T., Oba, Y., Takahashi, H., Kawai-Yamada, M., Uchimiya, H., Hashimoto, M., and Iba, K. (2008). CO2 regulator SLAC1 and its homologues are essential for anion homeostasis in plant cells. Nature 452:483–486.

Neher, E. (1992). Correction for Liquid Junction Potentials in Patch Clamp Experiments. Method Enzymol 207:123–131.

Qin, L., Tang, L.H., Xu, J.S., Zhang, X.H., Zhu, Y., Zhang, C.R., Wang, M.H., Liu, X.L., Li, F., Sun, F., et al. (2022). Cryo-EM structure and electrophysiological characterization of ALMT from Glycine max reveal a previously uncharacterized class of anion channels. Sci Adv 8:eabm3238.

Sasaki, T., Ariyoshi, M., Yamamoto, Y., and Mori, I.C. (2022). Functional roles of ALMT-type anion channels in malate-induced stomatal closure in tomato and Arabidopsis. Plant Cell Environ.

Sasaki, T., Mori, I.C., Furuichi, T., Munemasa, S., Toyooka, K., Matsuoka, K., Murata, Y., and Yamamoto, Y. (2010). Closing plant stomata requires a homolog of an aluminum-activated malate transporter. Plant Cell Physiol 51:354–365.

Schmidt, C., and Schroeder, J.I. (1994). Anion Selectivity of Slow Anion Channels in the Plasma Membrane of Guard Cells (Large Nitrate Permeability). Plant Physiol 106:383–391.

Schroeder, J.I., and Hagiwara, S. (1989). Cytosolic Calcium Regulates Ion Channels in the Plasma-Membrane of Vicia-Faba Guard-Cells. Nature 338:427–430.

Sievers, F., Wilm, A., Dineen, D., Gibson, T.J., Karplus, K., Li, W., Lopez, R., McWilliam, H., Remmert, M., Soding, J., et al. (2011). Fast, scalable generation of high-quality protein multiple sequence alignments using Clustal Omega. Mol Syst Biol 7:539.

Szyroki, A., Ivashikina, N., Dietrich, P., Roelfsema, M.R., Ache, P., Reintanz, B., Deeken, R., Godde, M., Felle, H., Steinmeyer, R., et al. (2001). KAT1 is not essential for stomatal opening. Proc Natl Acad Sci U S A 98:2917–2921.

Takanashi, K., Sasaki, T., Kan, T., Saida, Y., Sugiyama, A., Yamamoto, Y., and Yazaki, K. (2016). A Dicarboxylate Transporter, LjALMT4, Mainly Expressed in Nodules of Lotus japonicus. Mol Plant Microbe Interact 29:584–592.

Thomine, S., Guern, J., and Barbier-Brygoo, H. (1997). Voltage-dependent anion channel of Arabidopsis hypocotyls: nucleotide regulation and pharmacological properties. J Membr Biol 159:71–82.

Vahisalu, T., Kollist, H., Wang, Y.F., Nishimura, N., Chan, W.Y., Valerio, G., Lamminmaki, A., Brosche, M., Moldau, H., Desikan, R., et al. (2008). SLAC1 is required for plant guard cell S-type anion channel function in stomatal signalling. Nature 452:487–491.

Waterhouse, A.M., Procter, J.B., Martin, D.M., Clamp, M., and Barton, G.J. (2009). Jalview Version 2--a multiple sequence alignment editor and analysis workbench. Bioinformatics 25:1189–1191.

Zhang, J., Martinoia, E., and De Angeli, A. (2014). Cytosolic nucleotides block and regulate the Arabidopsis vacuolar anion channel AtALMT9. J Biol Chem 289:25581–25589.

