## Supplemental material for "ALMT-independent guard cell R-type anion currents"

**SUPPLEMENTAL MATERIAL (FIGURES, TABLES AND REFERENCES)**

**Fig. S1**

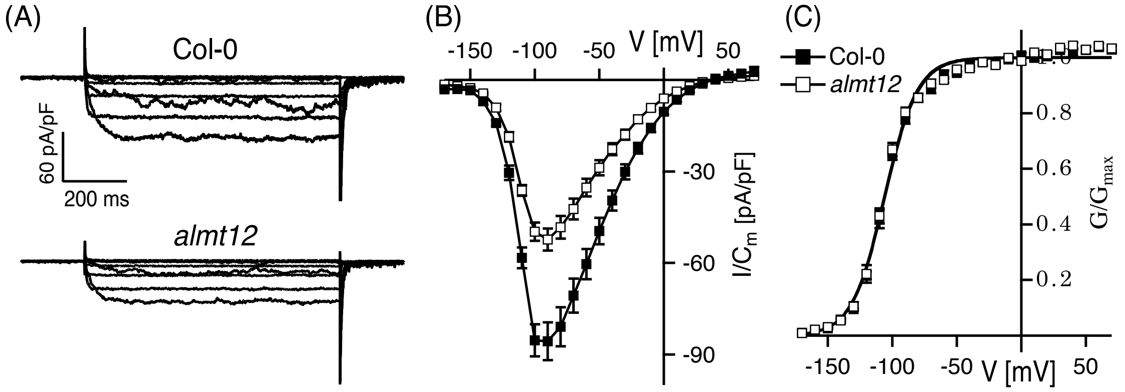


**Fig. S1. Voltage-dependent R-type current recordings from wildtype and *almt12* guard cell protoplasts.**

**(A)** Representative whole-cell currents shown at voltages in the range of -150 and +60 mV in 30-mV steps applied from a holding voltage of -170 mV.

**(B)** Normalized steady-state currents ($I/C_m$) plotted against membrane voltages applied from a holding voltage of -170 mV.

**(C)** Normalized conductances ($G/G_{max}$) plotted against membrane voltages. Solid line represents the best fit of the data points to a Boltzmann function. The derived midpoint voltages ($V_{1/2}$) and the apparent equivalent gating charges (z) were the following: $V_{1/2\text{-Col-0}}$ = -115 ± 1 mV, $V_{1/2\text{-}almt12}$ = -114 ± 1 mV, $z_{\text{Col-0}}$ = 2.02 ± 0.07, $z_{almt12}$ = 2.07 ± 0.06.

**(B, C)** Number of experiments were n=6 for Col-0 (closed squares), n=9 for *almt12* (open squares). Data points represent means ± SEM. Experiments in (**A-C**) were carried out with 195 nM free $Ca^{2+}$ and 5 mM ATP at the cytosolic side of the plasma membrane. Data presented in (**B**) are significantly different in peak current at -90 mV (***, p = 0.000293, One-Way Anova followed by Bonferroni test).

**Fig. S2**

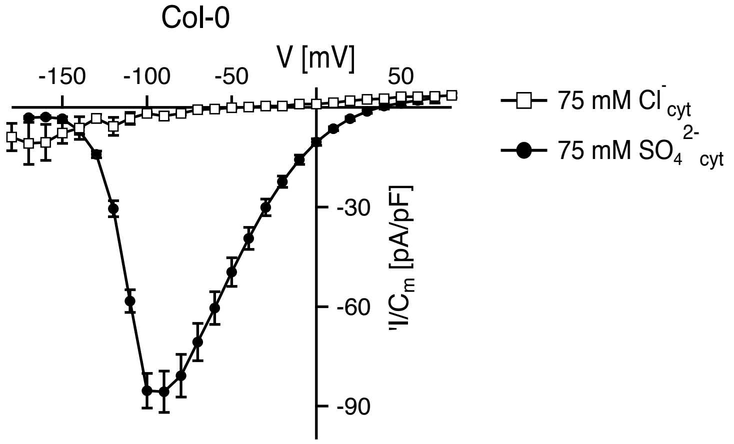


**Fig. S2. Permeation of wildtype R-type anion channels.**

Normalized whole-cell steady-state currents ($I/C_m$) of Col-0 guard cell protoplasts plotted against the membrane voltages. The holding voltage was -170 mV. Number of experiments were n=6 and n=3 for sulfate- and chloride-based pipette solutions, respectively. Data points represent means ± SEM. Experiments were performed in the presence of 195 nM free $Ca^{2+}$ and either 75 mM sulfate or chloride at the cytosolic side of the plasma membrane (cf. Imes et al., 2013).

**Fig. S3.**

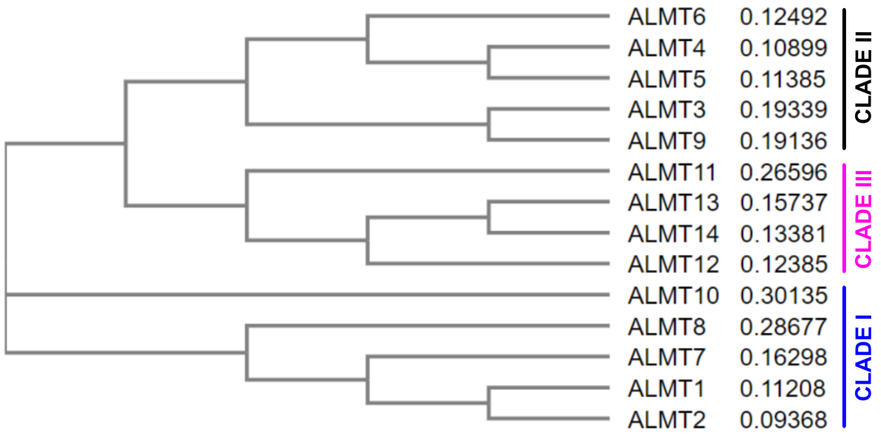


**Fig. S3. Dendrogram of the ALMT family**

Dendrogram of the *Arabidopsis thaliana* ALMT family divided into three clades (cf. Kovermann et al., 2007). Protein sequences were aligned using Hhalign algorithm. A guide tree was created using the neighbor-joining algorithm. After, each of the closet related pairs of sequences were aligned to each other. The phylogenetic tree was constructed using Clustalo Omega, version 1.2.4 (https://www.ebi.ac.uk/Tools/msa/clustalo/; Sievers et al., 2011) and visualized with Jalview 2.11.2.4 (Waterhouse et al., 2009). AtALMT1 (At1g08430), AtALMT2 (At1g08440), AtALMT3 (At1g18420), AtALMT4 (At1g25480), AtALMT5 (At1g68600), AtALMT6 (At2g17470), AtALMT7 (At2g27240), AtALMT8 (At3g11680), AtALMT9 (At3g18440), AtALMT10 (At4g00910), AtALMT11 (At4g17585), AtALMT12 (At4g17970), AtALMT13 (At5g46600), AtALMT14 (At5g46610).

**Fig. S4.**

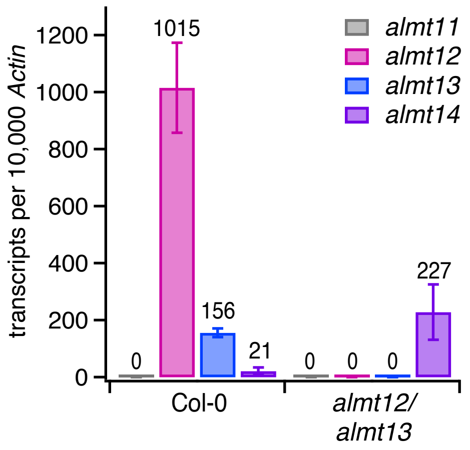


**Fig. S4. *ALMT* expression profile in *Arabidopsis thaliana* guard cell protoplasts of wildtype Col-0 and *almt12/almt13* mutant.**

Transcripts of *ALMTs* of clade III quantified relative to *ACTIN* transcripts. Expression level was quantified for guard cell protoplasts without pre-incubation in malate-containing solution. Data represent means ± SEM. Three biological repetitions were performed. Total number of examined samples was n = 10 for Col-0 and n = 8 for the *almt12/almt13* mutant.

**Fig S5.**

**A**

```
                            1                                                50
ALMT13 CDS WT               ATGGGGTACAAGGTCGAAGCACGGAGCATGGAGATTTCCATGGAAGATGA
ALMT13 CDS +G               ATGGGGTACAAGGTCGAAGCACGGAGCATGGAGATTTCCATGGAAGATGA
ALMT13 CDS -6bp             ATGGGGTACAAGGTCGAAGCACGGAGCATGGAGATTTCCATGGAAGATGA

                            51                                              100
ALMT13 CDS WT               AGATTCTAGAAAGAAGAGAAAGAAGGGTCTAAATCTTCCGAAGAAGATGA
ALMT13 CDS +G               AGATTCTAGAAAGAAGAGAAAGAAGGGTCTAAATCTTCCGAAGAAGATGA
ALMT13 CDS -6bp             AGATTCTAGAAAGAAGAGAAAGAAGGGTCTAAATCTTCCGAAGAAGATGA

                            101                                             150
ALMT13 CDS WT               AGAAGATTCTAAGGAATCTATGGAATGT-GGGAAAGGAAGATCCGAGGAG
ALMT13 CDS +G               AGAAGATTCTAAGGAATCTATGGAATGTgGGGAAAGGAAGATCCGAGGAG
ALMT13 CDS -6bp             AGAAGATTCTAAGGAATCTATGGA-------GAAAGGAAGATCCGAGGAG
```

**B**

```
                            1                                                50
ALMT14 CDS WT               ATGTCGGACAGGGTCCATGAAAGGAGCATGGGAATGGAGGAAGAAGGCTC
ALMT14 CDS +A               ATGTCGGACAGGGTCCATGAAAGGAGCATGGGAATGGAGGAAGAAGGCTC
ALMT14 CDS +T               ATGTCGGACAGGGTCCATGAAAGGAGCATGGGAATGGAGGAAGAAGGCTC
ALMT14 CDS 1SNP; -14bp      ATGTCGGACAGGGTCCATGAAAGGAGCATGGaAATGGAGGAAGAAGGCTC

                            51                                              100
ALMT14 CDS WT               TACTAAAAACATGAAG-A-CAAAGGTTCTTGAACTTCCAACAAAGATCAA
ALMT14 CDS +A               TACTAAAAACATGAAGaA-CAAAGGTTCTTGAACTTCCAACAAAGATCAA
ALMT14 CDS +T               TACTAAAAACATGAAG-AtCAAAGGTTCTTGAACTTCCAACAAAGATCAA
ALMT14 CDS 1SNP; -14bp      TAC----------------CAAAGGTTCTTGAACTTCCAACAAAGATCAA
```

**Fig S5. Multiple sequence alignment of *ALMT13* (A) and *ALMT14* (B) coding sequence obtained from wildtype (WT) Col-0 and *almt* triple lines.** CRISPR-Cas9-induced mutations are highlighted in turquoise background.

**Fig. S6.**

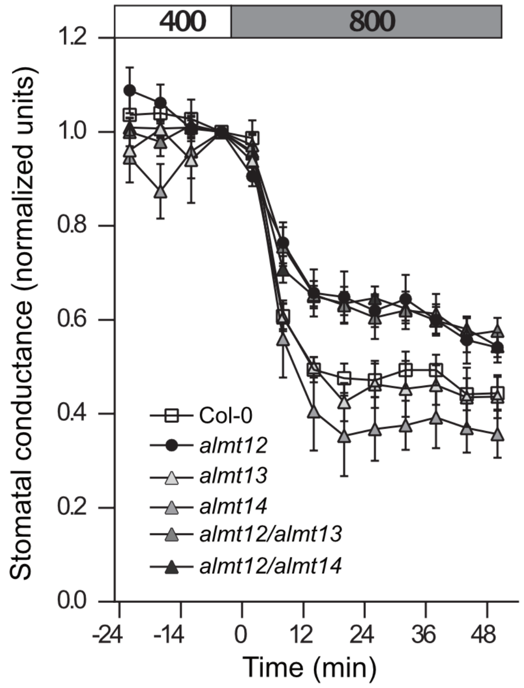


**Fig. S6. Gas exchange of different *almt12, almt13* and *almt14* mutant lines.** Normalized stomatal responses to changing $CO_2$ level from 400 ppm to 800 ppm at 0 time-point. Mean normalized stomatal conductance ± SEM is shown.

**Table S1. Values of half-activation voltages ($V_{1/2}$) and equivalent gating charges (z) determined for R-type anion conductance's of different ALMT loss-of-function mutants.**

| *A. thaliana almt* mutants | $V_{1/2}$ [mV] | z | n |
|---|---|---|---|
| *almt12* | -114.3 ± 1.3 | 2.07 ± 0.06 | 12 |
| *almt12/almt13* | -113.3 ± 0.8 | 1.94 ± 0.06 | 7 |
| *almt12/almt14* | -113.7 ± 1.5 | 1.99 ± 0.06 | 3 |
| *almt12/almt13/almt14#19* | -111.3 ± 0.7 | 2.79 ± 0.14 | 14 |

$V_{1/2}$ and z were derived from best fits of individual conductance voltage curves with a Boltzmann equation and are given as means ± SEM (n = number of individual experiments) (cf. Figs. 4B, 6D). Experiments were performed with 1.85 µM free $Ca^{2+}$ at the cytosolic side of the plasma membrane.

**Table S2. Primers used for generation of CRISPR-Cas9-based *almt* triple mutants.**

| Primer name | Sequence | Purpose |
|---|---|---|
| U6-26p-F | 5' TGTCCCAGGATTAGAATGATTAGGC 3' | Colony PCR primer of pHEE401E |
| U6-26t-R | 5' CCCCAGAAATTGAACGCCGAAGAAC 3' | Colony PCR primer of pHEE401E |
| ALMT12_F | 5' ACAAGACCACCGTTGGTAAACTC 3' | *almt12-2;* gene-specific genotyping |
| ALMT12_R | 5' CTCCGGCTAATCTTACACAAGG 3' | *almt12-2;* gene-specific genotyping |
| Spm32 | 5' TACGAATAAGAGCGTCCATTTTAGAGT 3' | *almt12-2;* T-DNA-specific genotyping |
| ALMT13_F1 | 5' CGCATTTCAATTATTAACGGCT 3' | Sequencing: *ALMT13* gDNA cut site |
| ALMT13_R1 | 5' ACTTACCGGCAGAGAACTCAAG 3' | Sequencing: *ALMT13* gDNA cut site |
| ALMT14_F1 | 5' CTTCGTCGGGCTAACTAGTCTC 3' | Sequencing: *ALMT14* gDNA cut site |
| ALMT14_R1 | 5' TAGGAATTTCTCGCAAATGGAT 3' | Sequencing: *ALMT14* gDNA cut site |
| ALMT14_F2 | 5' TCGGACAGGGTCCATGAAAG 3' | Sequencing: *ALMT14* gDNA cut site |
| ALMT14_R2 | 5' CCAAAGATCCAGCAATTAGTGTCC 3' | Sequencing: *ALMT14* gDNA cut site |
| ALMT13_F3 | 5' CGTAAGTAAACAACAAAACATAGAACAAG 3' | Sequencing: *ALMT13* cDNA |
| ALMT13_R4 | 5' ACTCACACATGCTTCAATACATCG 3' | Sequencing: *ALMT13* cDNA |
| ALMT13_F4 | 5' CACTTCTGTTTTCACCATCGGATCG 3' | Sequencing: *ALMT13* cDNA |
| ALMT13_R5 | 5' CACCTCGTGTTACAAAATTCCAG 3' | Sequencing: *ALMT13* cDNA |
| ALMT14_F1 | 5' CTTCGTCGGGCTAACTAGTCTC 3' | Sequencing: *ALMT14* cDNA |
| ALMT14_R4 | 5' CTCATTCACACATGCTTCGATGG 3' | Sequencing: *ALMT14* cDNA |
| ALMT14_F3 | 5' GCAGTTTTTATCATTGGAGCGTTG 3' | Sequencing: *ALMT14* cDNA |
| ALMT14_R5 | 5' CAAACCTCATCATACCTATTGGTTCG 3' | Sequencing: *ALMT14* cDNA |

**Table S3. Primers used for qRT PCR.**

| Gene | Forward Primer 5´→ 3´ | Reverse Primer 3´→ 5´ |
|---|---|---|
| ALMT1 | CTATACGAGAAGTCGGA | TGCCCATTACTTAATGT |
| ALMT2 | CCGTGGGTGCATTACT | TTCGACTTAGACGGCT |
| ALMT7 | CCTGTAAATCACTCACC | CGTCTACATGCGTATCA |
| ALMT8 | CACACATTGACAACTCC | ACATGACATGAGCTATT |
| ALMT10 | GATTTCCTGGTGTTTGA | GTTGTTTAGTGTTGTTGTC |
| ALMT11 | GACGATAAGCGAGAAAGTAAT | AGTATGGCTGGTTAAGGAC |
| ALMT12 | AGGAGGATGTTACGGC | GGTTTTCGCATCGGAC |
| ALMT13 | GGCCACGTTTCATATTAC | ACCTCCACAGTCTTATC |
| ALMT14 | AGGAAATAGCGGTAGAC | CTCAGAATCTTTCTTTCGT |
| ACT2/8 | GGTGATGGTGTGTCT | ACTGAGCACAATGTTAC |

### SUPPLEMENTAL REFERENCES

Imes, D., Mumm, P., Bohm, J., Al-Rasheid, K.A., Marten, I., Geiger, D., and Hedrich, R. (2013). Open stomata 1 (OST1) kinase controls R-type anion channel QUAC1 in Arabidopsis guard cells. Plant J 74:372-382.

Sievers, F., Wilm, A., Dineen, D., Gibson, T.J., Karplus, K., Li, W., Lopez, R., McWilliam, H., Remmert, M., Soding, J., et al. (2011). Fast, scalable generation of high-quality protein multiple sequence alignments using Clustal Omega. Mol Syst Biol 7:539.

Waterhouse, A.M., Procter, J.B., Martin, D.M., Clamp, M., and Barton, G.J. (2009). Jalview Version 2--a multiple sequence alignment editor and analysis workbench. Bioinformatics 25:1189-1191.
